# Ultra-sensitive coupling between organ growth and size by YAP-1 ensures uniform body plan proportions in *C. elegans*

**DOI:** 10.1101/2022.09.05.506666

**Authors:** Klement Stojanovski, Ioana Gheorghe, Anne Lanjuin, William B. Mair, Benjamin D. Towbin

## Abstract

Imbalance between the growth rate of different organs can amplify to large deviations of their size proportions during development. We show that, for the *C. elegans* pharynx, such size divergence is prevented by reciprocal coordination of pharyngeal growth with other tissues. Live imaging of hundreds of individuals revealed that small pharynxes grow more rapidly than large pharynxes, catching up in volume during development. Moreover, pharynx-to-body size proportions were robust to even strong tissue-specific inhibition of mTORC1 and insulin signalling. Tissue-specific depletion of these pathways slowed-down the growth of the respective tissue and additionally triggered a systemic growth response that ensured appropriate organ size proportions. By mathematical modelling, we show that the conservation of proportions requires a bi-directional ultra-sensitive coupling of body growth and pharynx size that cannot be explained by a reduction of food uptake alone. Instead, organ growth coordination requires regulation by the mechano-transducing transcriptional co-activator YAP/*yap-1*. Knock-down of *yap-1* makes animals sensitive to tissue-specific inhibition mTORC1 inhibition, causing a disproportionate pharynx and developmental arrest. Our data suggests that mechano-transduction tightly coordinates organ growth during *C. elegans* development to ensure the uniformity of body plan proportions among individuals.

## Introduction

The correct size and proportions of organs are crucial for organismal function, and disproportionate organs are associated with a large diversity of diseases ^1–4^. However, attaining correct size proportions is challenged by fluctuations in organ growth rates. Since organs grow by orders of magnitude during development, in principle, even small deviations from the correct organ growth rates could amplify to large deviations in their size over time ^5,6^. Size divergence of organs is usually not observed in nature, but the mechanisms that prevent divergence are not well understood.

Size homeostasis has been studied extensively at the scale of individual cells. Quantitative time-lapse microscopy of yeasts, bacteria, and mammalian cells revealed that stochastic fluctuations of cell size are corrected within a few cell divisions ^7–11^. Cell size correction often involves a so-called adder or sizer mechanisms^7–11^, in which large cells undergo a smaller volume fold change per cell cycle than small cells. These cell autonomous mechanisms break the intrinsic tendency of exponential growth to amplify differences in cell size, such that cells that deviate from the norm return to a stable reference point within a few cell cycles. We have recently shown that, for *C. elegans*, size uniformity at the scale of the entire body volume does not follow adders or sizers ^12^. Instead, the volume fold change per larval stage is near independent of the starting size. Nevertheless, rapidly and slowly growing individuals only weakly diverge in volume due to an inverse coupling of the growth rate to the duration of development by a genetic oscillator, a mechanism we termed a growth-coupled folder ^12^.

At the scale of organs, size control involves tissue-autonomous and systemic mechanisms. For example, morphogen gradients are thought to limit the lateral expansion of imaginal discs in *Drosophila melanogaster* and thereby limit their final size tissue autonomously ^13,14^. At the same time, experiments using *Drosophila* also provide evidence for systemic size control mechanisms ^15,16^. For example, growth inhibition of the *Drosophila* wing disc triggers the release of the relaxin-like signalling peptide *Dilp8*, which systemically modulates larval growth through the growth hormone ecdysone ^17,18^. Similarly, unilateral inhibition of limb growth in mice triggers a growth response that retains proper limb symmetry ^19,20^. Despite these important advances, it remains unclear how the many tissues of an organism coordinate their growth to robustly reach the correct size proportions.

Here, we used the pharynx of *C. elegans* to ask if the organ size uniformity involves adders and sizers, and whether this control is systemic or autonomous. We find that, similar to adder-like mechanisms described in unicellular organisms, spontaneously occurring deviations in pharynx size were corrected by size-dependent growth regulation. However, in contrast to bacteria, size control was not cell autonomous and involved crosstalk between different cells: inhibition of growth signalling in the pharynx triggered a systemic response in other tissues and vice versa. This bi-directional growth response is ultra-sensitive to deviations in pharyngeal size of a few percent and cannot be explained by limitation of food uptake alone. Consequently, the scaling between pharynx and body size is nearly invariant even under strong tissue-specific growth perturbations. Genetic experiments revealed that the mechano-transducing transcriptional coactivator *Yes associated protein* (YAP)^21^ is required for organ growth coordination, suggesting an important role of mechano-sensing in the robustness of organ size proportions of *C. elegans*.

## Results

### Quantification of pharyngeal and total body growth of *C. elegans* by live imaging

To quantify the growth and size of the pharynx relative to other tissues, we created a *C. elegans* strain expressing a green fluorescent protein in the pharyngeal muscle (*myo-2p::gfp*), and ubiquitously expressing a red fluorescent protein (*eft-3p:mscarlet*) (Fig. 1a). We recorded growth of hundreds of individual animals of this strain at 25 °C in arrayed micro chambers using a temperature-controlled fluorescence microscope at a time resolution of 10 minutes ^12^. By automated image analysis ^12^, we determined the length of the pharynx and the total body (Supplemental Fig. 1a) at each timepoint from planar optical sections, and inferred their volumes given their near rotational symmetry (Fig. 1b) ^22^.

**Figure 1.**
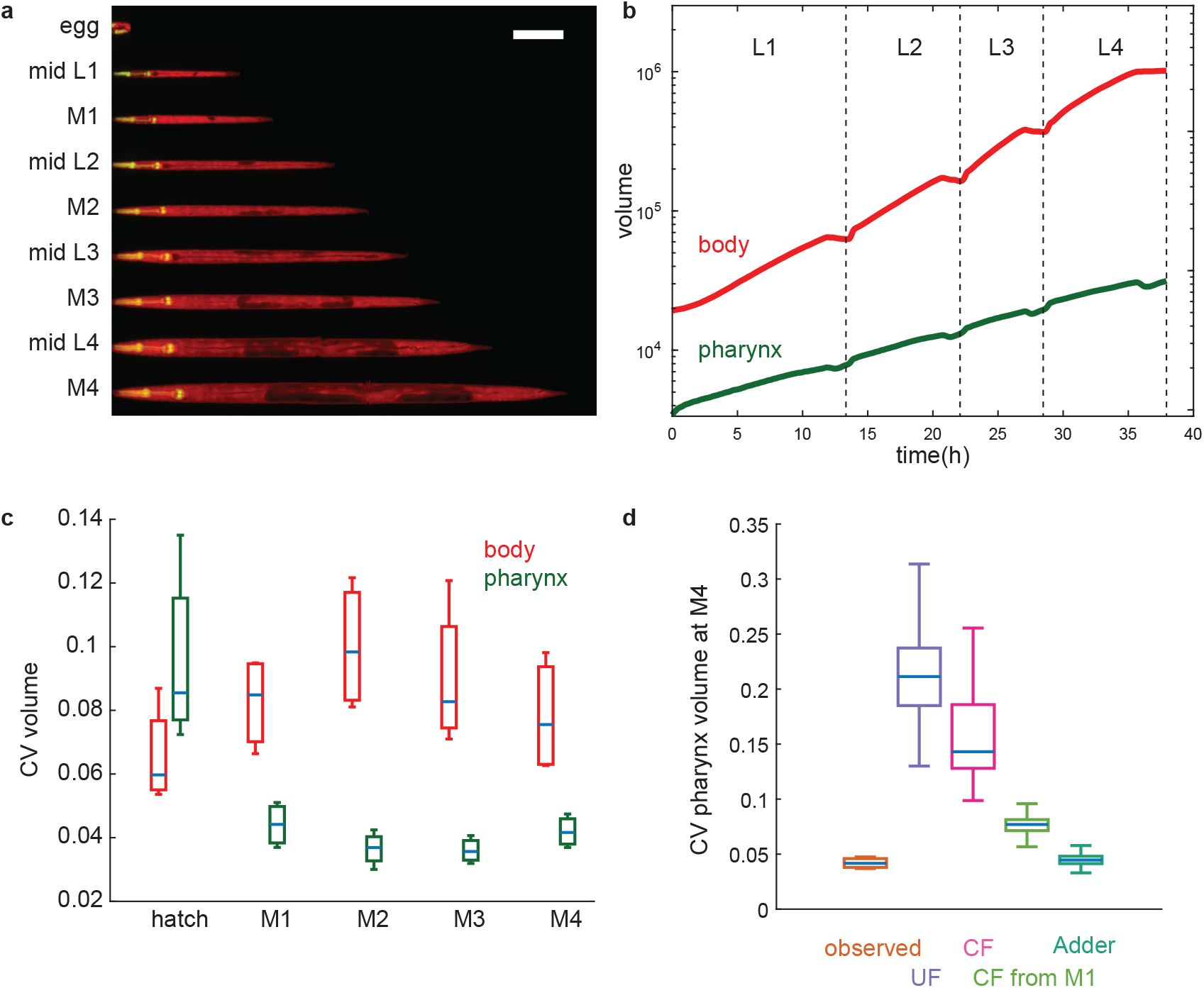
Pharyngeal volume is less heterogeneous among individuals than total body volume. a. Individual animal imaged in micro chambers at indicated developmental milestones. Merged image of pharynx in green (*myo-2p:gfp*) and total body in red (*eft-3p:mscarlet*) is shown. Overlay of green and red is shown as yellow. *eft-3p:mscarlet* is comparatively weak in the head region in young L1 animals, producing a more green appearance of the pharynx at this stage. Contrast was adjusted for each time point individually. Animals were straightened computationally. Scale bar: 100μm b. Body (red) and pharynx (green) volume as a function of time averaged from n = 475 individuals. For averaging, individuals of each larval stage were re-scaled to have matching larval stage entry and exit points and the growth curve was scaled back to the mean larval stage duration after averaging. c. Coefficient of variation (CV) of body (red) and pharynx (green) volume at hatch and larval moults (M1 to M4). Box plots represent CVs of 4 independent day-to-day repeats with 105 < n < 181 individuals per repeat. d. Comparison of observed CV of pharyngeal volume to randomized simulations as described in the main text. UF: uncoupled folder, CF: coupled folder, CF from M1: CF starting at M1. p-value (one-sided t-test) for observed vs. model: UF: 0.0007, CF: 0.0083, CF from M1: 10^−5^, Adder: 0.04. Boxplots in c,d: central line: median, box: interquartile ranges (IQR), whisker: ranges except extreme outliers (>1.5*IQR)

As we previously observed ^12^, total body growth was halted during four periods of approximately two hours, corresponding to lethargic phases prior to cuticular moulting (Fig. 1b). We could thereby automatically detect larval stage transitions by the time points at which growth re-started and feeding resumed and compute growth rates μ of body and pharynx at each larval stage as the increase in log(volume) per time. Throughout this article growth rate refers to the change in log transformed volume per time, i.e. the growth rate normalized to the current size, unless specified as the absolute growth rate μ_abs_, which indicates the absolute change in volume per time.

The average growth rate of the pharynx was about half the growth rate of the total body volume (doubling times of 12.02 +/- 0.94 vs. 6.64 +/- 0.51 hours). Consistently, the cumulative volume fold change from hatch to the fourth moult (M4) was 6 times larger for the total body (53.3 +/- 5.3 fold) than for the pharynx (9.02 +/- 1.05 fold). Similar to allometric growth of the head and body of many animals, the volume fraction of the pharynx thus declined from 18 +/- 1.7% at hatching to 3.1 +/- 0.27 % at M4.

### Pharyngeal size heterogeneity does not increase during development

During development, small differences in the growth rates among individuals are prone to amplify to large differences in size, particularly during exponential growth. As we have previously shown, the coefficient of variation (CV) of the body volume among individuals nevertheless remains below 10% ^12^ (CV of body length: ^~^4%) (Fig. 1c, Supplemental Fig. 1b). This maintenance of body size uniformity occurs largely due to an inverse coupling of growth to the duration of larval stages, and not due to sizers or adders ^12^.

Remarkably, the CV of the pharyngeal volume (4%) and length (2%) was even smaller than for total body size (Fig. 1c, Supplemental Fig. 1c), except for the size at hatching. To ask if the degree of size heterogeneity was smaller than expected given the heterogeneity of the pharyngeal growth rates, we ran simulations in which we randomly shuffled measured growth parameters among individuals. We then compared the measured size heterogeneity to those produced by randomized simulations. Building on our previous work^12^, our simulations distinguished four different scenarios. First, we independently randomized the growth rates and larval stage durations among individuals (a model called *uncoupled folder*). Second, we randomly shuffled the volume fold changes per larval stage among individuals but retained the coupling between growth and larval stage duration (*coupled folder*). Third, we simulated a coupled folder, but started simulations at the first moult (M1) instead of at hatch. This comparison is motivated by our previous description of unique features to size control at the L1 stage ^12^. Fourth, we simulated an adder mechanism by randomly shuffling the added volume per larval stage among individuals. Out of these four simulations, only the simulated adder was close to the observed heterogeneity. Thus, unlike for total body volume ^12^, a coupled folder cannot recapitulate the smaller size heterogeneity of the pharynx. Together, these data suggest that the pharynx is under more stringent size control than the total body volume (Fig. 1d, Supplemental Fig. 1c) ^12^.

### Sub-exponential pharyngeal growth within one larval stage results in an adder-like behaviour

Under exponential growth and in the absence of size-dependent control of growth, the volume growth per time or developmental stage is expected to scale with the current size of an individual. Indeed, the added total body volume per larval stage of *C. elegans* was positively correlated with the volume at larval stage entry (Fig. 2a, Supplemental Fig. 2b), except for L1 animals as we previously described ^12^. However, for the pharynx, we did not observe this same relation. Instead, the absolute volume increase ΔV during L2 and L4 stages was nearly independent of the starting volume V_1_, reminiscent of an adder mechanism ^7,9,10^. During L1 and L3, we observed an even more stringent size-dependent control of pharynx growth, as ΔV and V_1_ were anti-correlated in these stages (Fig. 2b, Supplemental Fig. 2a).

**Figure 2.**
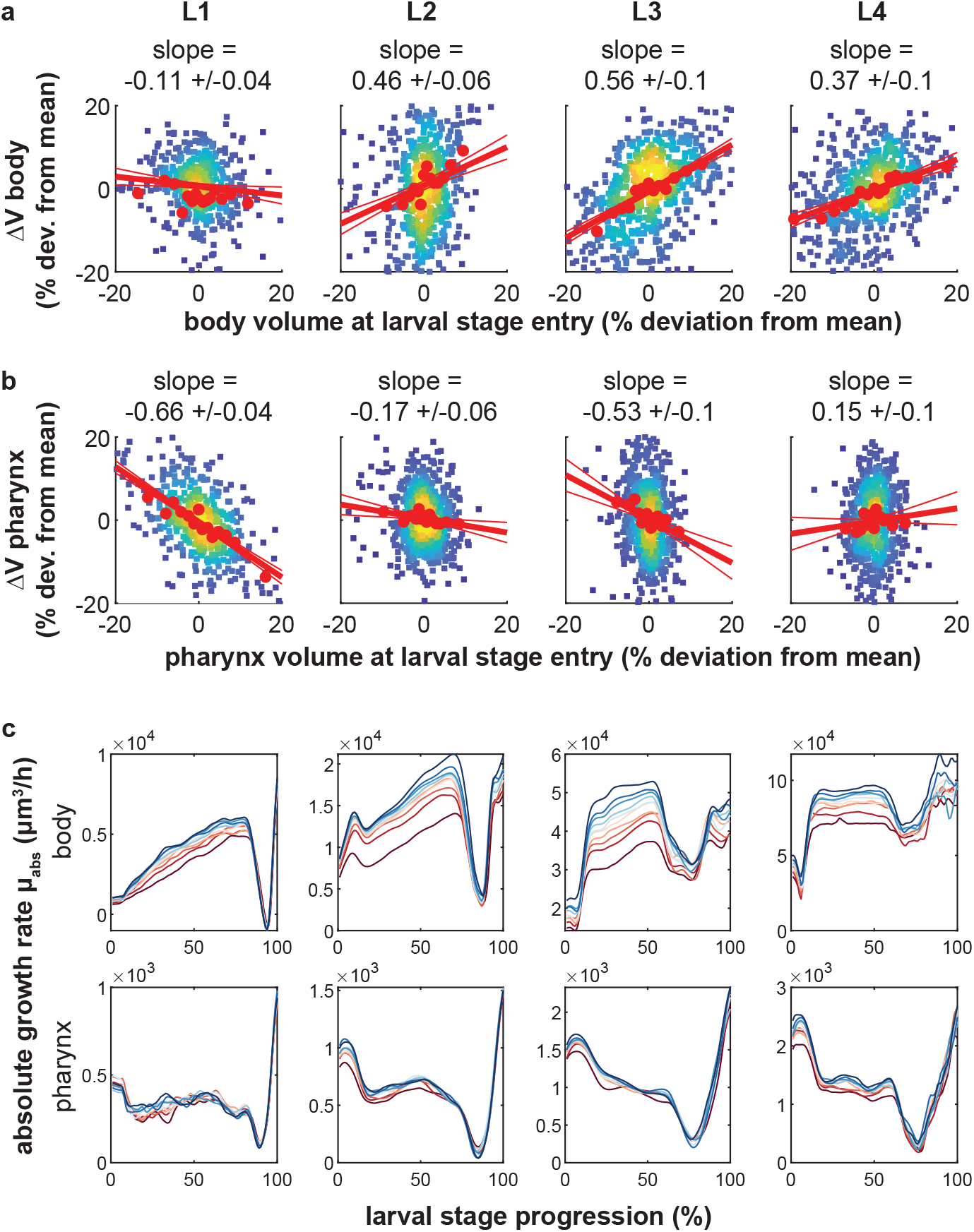
Linear pharyngeal volume growth within larval stages produces and adder-like behaviour. a. Scatter plot of body volume at larval stage entry vs. added body volume per larval stage for each individual. Red circles: binned average along x-Axis. red line: robust linear regression (thick) with 95% confidence intervals (thin). Data is shown as relative deviation to the batch mean. Rare outliers beyond the +/- 20% range are omitted for clarity. Value above chart indicates slope of regression line +/- 95% CI b. As a., but for pharyngeal volume. c. Absolute rate of body (top) and pharynx (bottom) volume increase as a function of larval stage progression. Individuals were binned into 10 classes according to their volume at 40% of each stage (red is the smallest volume, blue the largest volume). Individuals were re-scaled from moult to moult before averaging. The drop in growth rate at 80-90% of the larval stage represents growth halt during lethargus. a-c. n = 475,641, 641, 638 individuals for L1 to L4 from 4 day-to-day repeats.

To study the dynamics of pharyngeal size control, we analysed the volume growth rate trajectories within each larval stage for animals of different size. We binned individuals according to their pharynx or body volume in the middle of each larval stage and calculated the absolute rate of volume increase (μ_abs_ = dV/dt) as a function of larval stage progression. Consistent with autocatalytic and supra-linear body growth^12^, μ_abs_ was correlated with the body size and was larger at the end than at the beginning of each larval stage (Fig. 2c). However, although μ_abs_ of the pharynx increased near exponentially between larval stages (Supplemental Fig. 2d), μ_abs_ was nearly constant within a given larval stage (Fig. 2c). Moreover, large and small pharynxes had nearly the same μ_abs_ (Fig. 2c, Supplemental Fig. 2d). We conclude that although pharyngeal size trajectories closely mimic exponential growth when inspected across multiple larval stages (Fig. 1b, Supplemental Fig. 2d), the pharynx grows near linearly over time within each larval stage. Linear growth at an absolute growth rate μ_*abs*_ that is independent of the starting size is expected to produce a size-independent volume increase Δ*V* per time (*V*(*t*) = *V*_0_ + μ_*abs*_ * *t* = *V*_0_, + Δ*V*), which is consistent with the observed adder mechanism of the pharynx.

As described above, simulations of an adder can indeed reproduce the observed coefficient of variation in pharyngeal volume, whereas other models produce a size divergence that is larger than we experimentally observed (Fig. 1d, Supplemental Fig. 1c). Notably, we do not find a correlation between the pharynx size and the body growth rate among individuals, suggesting that the pharynx size is not limiting for growth in the observed range, and that uniformity of pharynx size is not a passive consequence of changes in the food uptake rate (Supplemental Fig. 2e).

### Auxin-inducible degradation of RagA/RAGA-1 allows quantitative titration of growth rates

Size control of the pharynx could be tissue-autonomous or involve crosstalk with other tissues. To distinguish between these two scenarios, we developed an experimental approach to perturb growth tissue-specifically by auxin-induced degradation (AID) ^23^ of the mTORC1 activator RagA/RAGA-1 ^24–26^. To this end, we expressed the plant ubiquitin ligase Tir1 under the control of ubiquitous or tissue-specific promoters and modified the endogenous locus of *raga-1* with an aid-tag. Supplementation of the plant hormone auxin (indole-3-acetic-acid, IAA) leads to dosage dependent ubiquitination of aid-tagged proteins and consequently their quantitative knock-down by proteasomal degradation in the tissue where Tir1 is expressed ^27^. We could thereby perturb growth by depleting RAGA-1 by AID (Fig. 3a) and measure length and volume growth in micro chambers. Length and volume measurements led to equivalent conclusion. In the following, for their higher accuracy, length measurements are shown in the main figures. The corresponding volume data is displayed in Supplemental Figures.

**Figure 3.**
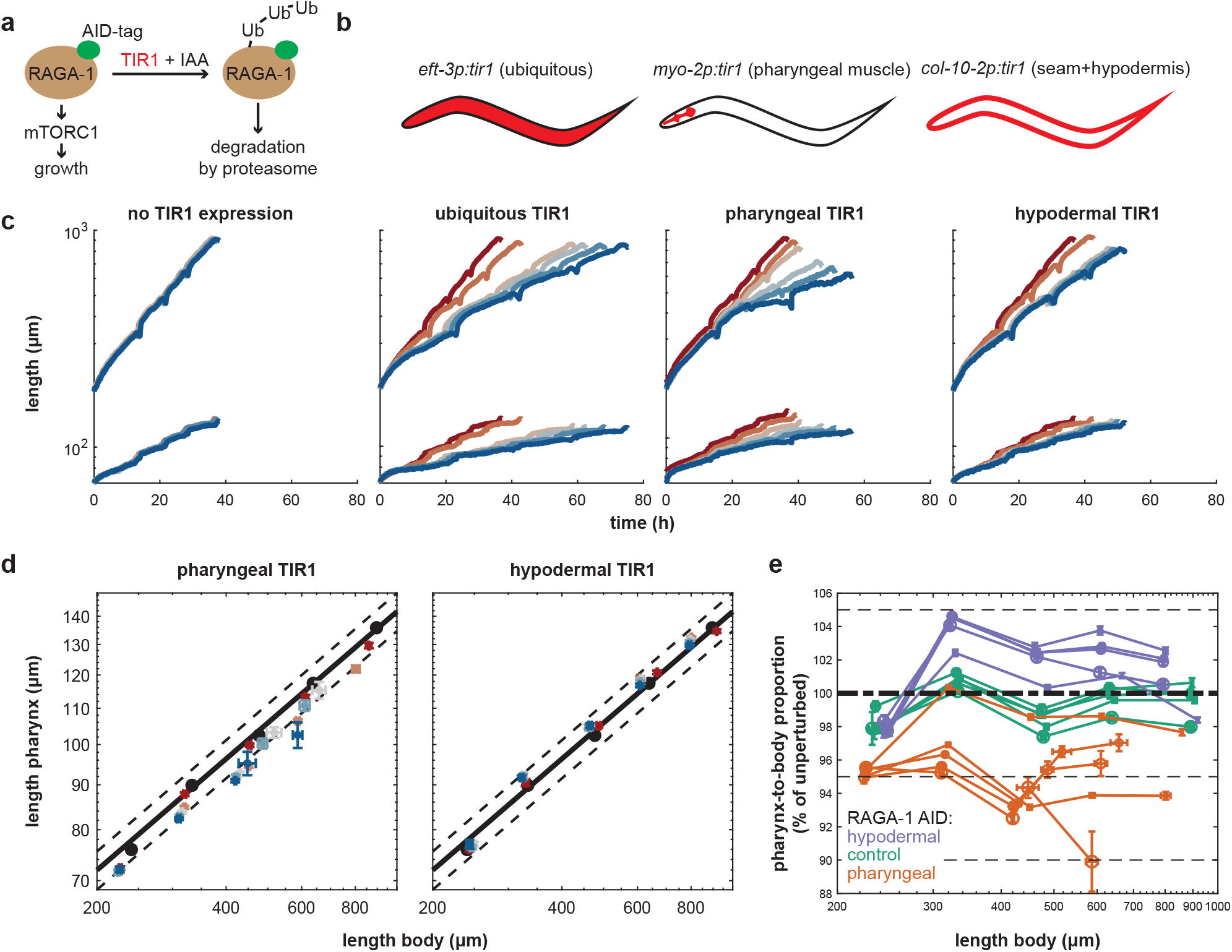
Pharynx-to-body length proportions are robust to tissue-specific depletion of RAGA-1. a. Experimental approach to deplete RAGA-1 in selected tissues. An AID tag was inserted into the endogenous locus of the *raga-1* gene and Tir1 was expressed ubiquitously or under tissue-specific promoters. Addition of auxin to the growth medium leads to ubiquitination and proteasomal degradation of RAGA-1 in tissues expressing Tir1. b. Schematic representation of tissue-specific promoters used to express the Tir1 enzyme. c. Body (top) and pharynx (bottom) length as a function time with depletion of RAGA-1 in indicated tissues. Color indicates IAA concentration from red to blue: no Tir1 expression 0mM IAA, Tir1 expression + 0mM, 0.1mM, 0.25mM, 0.5mM, 1mM IAA. Number individuals n, 26 < n < 259 from number of day-to-day repeats m, 3 < m < 11. See Supplemental Tables 1 and 2 for n and m of each condition. d. Scatter plot showing body vs. pharynx length at the beginning of L1 and at all larval moults M1 to M4 (circles in order from left to right) under pharyngeal (left) or epidermal (right) AID of RAGA-1. black circles: relation between pharynx and body length when unperturbed (no Tir1 expression and no IAA). Solid black line: linear regression to unperturbed body-to-pharynx length (P-line). Dashed black line: 5% deviation from P-line. Coloured circles: IAA concentrations from red to blue: 0, 100, 250, 500, 1000μM. Error bars are standard error of the mean among day-to-day repeats. e. Deviation of pharynx-to-body length ratio (deviation from P-line) vs. body length at early L1 and larval moults. Colours indicate different Tir1 expression; orange: pharynx, purple: epidermis, green: no Tir1 expression. Circle size indicates IAA concentration from 0mM (smallest circle) to 1mM (largest circle) as indicated in d. Error bars are standard errors among day-to-day repeats.

Consistent with the effect of *raga-1* deletion mutants^12^, ubiquitous AID of RAGA-1 reduced pharynx and body growth rates and extended larval stage durations (Fig. 3c, Supplemental Fig. 3a-b). Growth inhibition was quantitatively dependent on the strength of knock-down and scaled with the IAA concentration applied. We note that Tir1 has weak activity towards the AID tag even in the absence of IAA ^28,29^, such that growth was reduced by 10-21% (depending on the larval stage) compared to a strain not expressing Tir1. Throughout this study, we therefore include a strain lacking Tir1 expression, in addition to omitting IAA as a negative control.

### Pharynx specific inhibition of RagA/*raga-1* reduces growth, but retains pharynx-to-body size proportions

To ask how tissue-specific AID of RAGA-1 affects the allometric relation between the pharynx and the rest of the body, we used two strains expressing Tir1 only in selected tissues ^23,27^ (Fig. 3b). First, we expressed Tir1 specifically in the pharyngeal muscle under the control of the *myo-2* promoter ^23^. Second, we restricted Tir1 expression to hypodermal and seam cells ^27^ (collectively referred to as epidermis ^30^ from here onwards) using the *col-10* promoter. In either of these strains, pharyngeal as well as body growth were quantitatively reduced in an IAA dependent manner (Fig. 3c, Supplemental Fig. 3a-b). Thus, depletion of RAGA-1 in a single tissue reduces growth non-autonomously also in other tissues. Epidermal RAGA-1 AID reduced growth less strongly than pharyngeal RAGA-1 AID, which may be due to biological or technical differences in the effectiveness of reducing RAGA-1 or mTORC1 activity in these two tissues.

To quantify the non-autonomous response between pharynx and body growth, we plotted log-transformed length of body and pharynx at all four moults (M1 to M4) against each other (Fig. 3d). An equivalent analysis of volumes is shown in Supplemental materials (Supplemental Fig. 3c-d). After log transformation, body and pharynx length were near linearly related, as is expected for two tissues with near exponential growth. We call this relationship the P-line (P for pharynx), where the slope 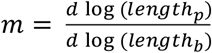 of the P-line corresponds to the ratio of the exponential growth rate of pharynx and body 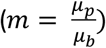. Analysis of the P-line under perturbed conditions thus allows us to distinguish different scenarios: (i) In the absence of tissue growth coordination, tissue-specific inhibition of growth is expected to change the ratio between pharynx and body growth rates and thus alter the slope *m* of the P-line. (ii) If unperturbed tissues respond with a proportional change in their growth rate the slope of the P-line remains constant. (iii) A temporary deviation from the unperturbed organ growth proportions early in development, followed by an appropriate adjustment of systemic growth rates would produce a P-line with unchanged slope, but a change in y-intercept.

We note that in some experiments, we observed a very rapid length extension immediately after hatching (Fig. 3c). This effect is likely of technical origin and may relate to a weak penetration of IAA through the egg shell. To avoid confounding effects, we therefore start our analysis at 30% passage of the L1 stage rather than immediately after hatching. Epidermal as well as pharyngeal RAGA-1 AID were in close agreement with scenario (iii) described above. Under pharyngeal RAGA-1 AID, the P-line was shifted down by 5-8% but did not systematically change in slope (Fig. 3d). This was true even at very high IAA concentrations and throughout development, except for the highest IAA concentration at the last moult. Similarly, epidermal RAGA-1 AID caused a near parallel upshift of the P-line by less than 5% without systematically changing its slope. Thus, in both tissues, RAGA-1 AID resulted in only a small difference in length proportions that plateaued at 5-8% and that did not scale with the degree of RAGA-1 inhibition (Fig. 3e, Supplemental Fig 3e). This saturation of pharyngeal length deviation suggests that tissues in which RAGA-1 was not depleted by AID reduced their growth rate near proportionally to the tissue targeted by AID. Pharynx-to-body length proportions thereby remained nearly constant even under strong tissue-specific impairment of mTORC1 signalling. We note that while pharynx-to-body proportions are retained (Fig. 3d,e), pharyngeal depletion of RAGA-1 results in a strong reduction of overall size (Fig. 3c), indicating that body size and pharynx-to-body proportions are controlled by separable mechanisms.

Repeating the same experiments using an AID-tagged allele of the insulin signalling receptor DAF-2 instead of RAGA-1 produced near identical results. Like AID of RAGA-1, pharyngeal size deviation plateaued at less than 5% deviation despite significant growth inhibition of the pharynx (Supplemental Fig. 4). The observed robustness in pharynx length proportions is thus not a specific response to RAGA-1 inhibition, but a more general response to an imbalance in tissue growth or size. Analysis of volume instead of length led to equivalent conclusions (Supplemental Fig. 3c), with only minor quantitative differences. For pharyngeal depletion of RAGA-1, the maximal deviation of the pharyngeal volume was larger than for length (15% vs. 5%). For epidermal RAGA-1 AID, volume deviations were smaller than length deviations (−3% vs. +5%, Supplemental Fig. 3d). This difference between the effects on length and volume is explained by slightly altered body length-to-width proportions under epidermal RAGA-1 AID (Supplemental Fig. 3e).

We conclude that pharynx-to-body size proportions are highly robust to even strong tissue-specific inhibition of growth.

### Body growth has ultra-sensitive dependence on pharynx size

To quantify the mutual relation between size and growth of pharynx and body, and to estimate how deviations from this relation would affect pharynx-to-body proportions, we compared the experimental data to the following mathematical model (Fig. 4b).

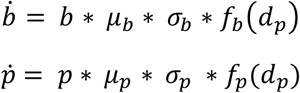

μ_p_ and μ_b_ are the growth rates of pharynx size *p* and body size *b* in the absence of experimental perturbation. σ_p_ and σ_b_ indicate the degree of experimental growth inhibition (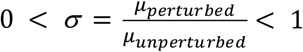, i.e. *σ* = 1 corresponds to unperturbed growth). 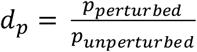 corresponds to the deviation from the unperturbed pharynx size proportions. *f_b_* and *f_p_* describe how body and pharynx growth relate to the pharyngeal size deviation *d_p_*.

**Figure 4.**
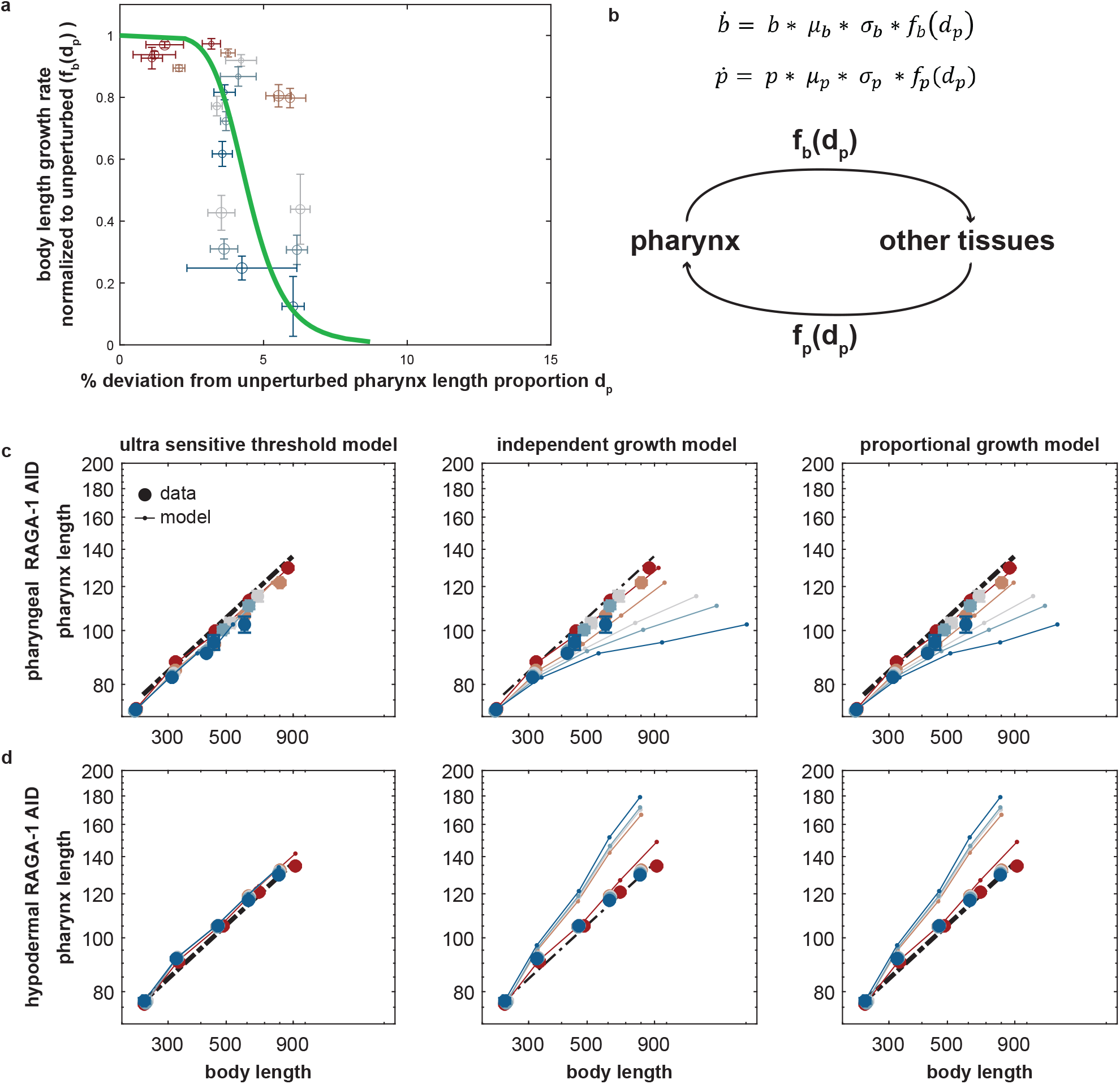
An ultra-sensitive relation between pharynx size and body growth is required for robustness of pharynx-to-body length proportions to tissue-specific growth inhibition. a. Body length growth rate (Δlog(len)/Δt) as a function of deviation of pharynx length dp from the P-line under pharyngeal growth inhibition. Growth is normalized to unperturbed growth. Colour indicates IAA concentration increasing from red (0 mM) to blue (1 mM). Circle size indicates the larval stage (L1 being the smallest). Green line is a fitted Hill function as described in the main text. b. Schematic illustration of mathematical model as described in the text. c. Comparison of experimental data (solid circles) from pharyngeal RAGA-1 AID to three different models (thin lines) as described in the main text with 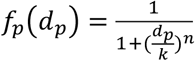 (left), *f*(*d_p_*) = 1 (middle), *f*(*d_p_*) = 1 – *d_p_* (right). Colours as in a. The ultra-sensitive threshold model, but not proportional or the independent model fit the data. d. As c., but for epidermal RAGA-1 AID. a,c,d. error bars in vertical and horizontal direction indicate standard error of the mean between day-to-day repeats. Absence of error bars means that they were smaller than the marker size.

Experimentally, *f_b_*(*d_p_*) and *f_p_*(*d_p_*) can be determined by plotting the measured deviation from the unperturbed pharynx size *d_p_* against the observed reduction in body or pharynx growth rates (Fig. 4a). This analysis showed that *f_b_*(*d_p_*) is a highly non-linear function approximating the Hill function 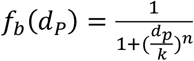 with half-inhibition constant k=4.4% and a Hill coefficient n=6.8.

Thus, we find that the measured body length growth rates rapidly drop at even very small deviations from pharynx length: body growth is reduced by more than 50% at less than 5% deviation of the pharynx length. The reverse relation 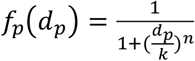 had a similar half-way point of about 5%, although the available experimental data did not allow a quantitative fit to the model (Supplemental Fig. 5a). Like lengths, the relation between pharynx volume and the body growth rate is also well approximated by a Hill function (Supplemental Fig. 5b), albeit with slightly different parameters. In conclusion, under tissue-specific depletion of RAGA-1, the body growth rate has ultra-sensitive dependence on the deviations from the appropriate pharynx size and vice versa.

### Ultra-sensitive coupling of body and pharynx growth and size ensures correct pharynx size proportions

To ask if the ultra-sensitive relation between pharynx and body growth is required for the observed robustness of pharynx size proportions, we simulated pharynx and body growth by the above-described model using three different expressions for *f_b_*(*d_p_*) and *f_p_*(*d_p_*). First, we used the fitted ultra-sensitive Hill function (see above). Second, we simulated independent growth of pharynx and body (*f*(*d_p_*) = 1), and third we modelled proportional scaling of body growth to pharynx size (*f*(*d_p_*) = 1 – *d_p_*). Proportional scaling approximates an effect expected by a proportional limitation of food uptake due to a smaller pharynx.

Simulations using the measured Hill function produced good fit to the experimental data. This agreement between data and model suggests that a single Hill function *f*(*d_p_*) = *f_p_*(*d_p_*) = *f_b_*(*d_p_*) for all developmental stages, IAA concentrations, and tissues is sufficient to accurately describe the observed growth dynamics. Thus, stage- and condition-specific deviations from the idealized Hill function (Fig. 4a, Supplemental Fig. 5a) do not substantially contribute to the observed invariance of pharynx-to-body size proportions. Contrastingly, the two alternative models strongly deviated from the experimental observations (Fig. 4c,d). We conclude from this comparison between data and model that proportional dependence between pharynx size and growth is insufficient to explain the experimental observations. Instead, an ultra-sensitive relation between pharynx size and body growth is needed to produce the observed robustness in pharynx size proportions.

### YAP-1 is required for the robustness of pharyngeal size proportions to epidermal growth inhibition

The ultra-sensitive dependence between pharynx size and body growth suggested that pharynx-to-body length coordination involves molecular regulatory mechanisms and cannot be explained by an indirect effect of restricted nutrient uptake. To identify molecules involved, we used RNAi and genetic mutations to systematically impair canonical growth regulatory pathways. Specifically, we targeted genes associated with (i) transforming growth factor β (TGFβ) ^31^, (ii) insulin and insulin-like growth factor (IIS) ^32^, (iii) mTOR ^33^, and (iv) Yes-associated protein/YAP signalling ^34^. For each gene, we exposed animals with a *raga-1-aid* tag expressing Tir1 in the epidermis or in the pharynx to RNAi and imaged growth of individual animals in micro chambers. In parallel, we used a strain lacking Tir1 expression as a control for each RNAi to distinguish genes involved in a response to imbalanced tissue growth from those that are more generally involved in controlling pharynx size.

Our systematic analysis revealed the gene *yap-1* as essential for retaining pharynx-to-body proportions when RAGA-1 was depleted selectively in the epidermis (Fig. 5a,b). YAP-1 is the *C. elegans* ortholog of the human transcriptional co-activator YAP ^34^, which functions as a mechano-transducer downstream of the hippo signalling pathway ^35^. Notably, the effect on organ size proportion was specific for *yap-1* RNAi and was not observed for other genes, although we cannot exclude insufficiency of knock-down for this lack of effect (Supplemental Fig. 6). For example, we did not detect synergistic effects between RAGA-1 AID and RNAi of the TGFβ ligand *dbl-1* or its downstream effector gene *lon-1* (Fig. 5a,b and Supplemental Fig. 6b). Similarly, a null mutation of the IIS effector gene *daf-16* did not perturb pharynx-to-body growth coordination. Although *daf-16* mutants had a slightly reduced pharynx length under otherwise unperturbed conditions, this difference was not enhanced by tissue-specific RAGA-1 AID (Supplemental Fig. 6a). Similarly, we did not identify any RNAi that enhanced the sensitivity of pharynx-size proportions to pharyngeal RAGA-1 AID (Supplemental Fig. 6a). This higher sensitivity of *yap-1(RNAi)* animals to epidermal over pharyngeal AID of RAGA-1 suggests tissue-specificity in the mechanisms of organ growth coordination, although we cannot exclude that this difference is of technical origin.

**Figure 5.**
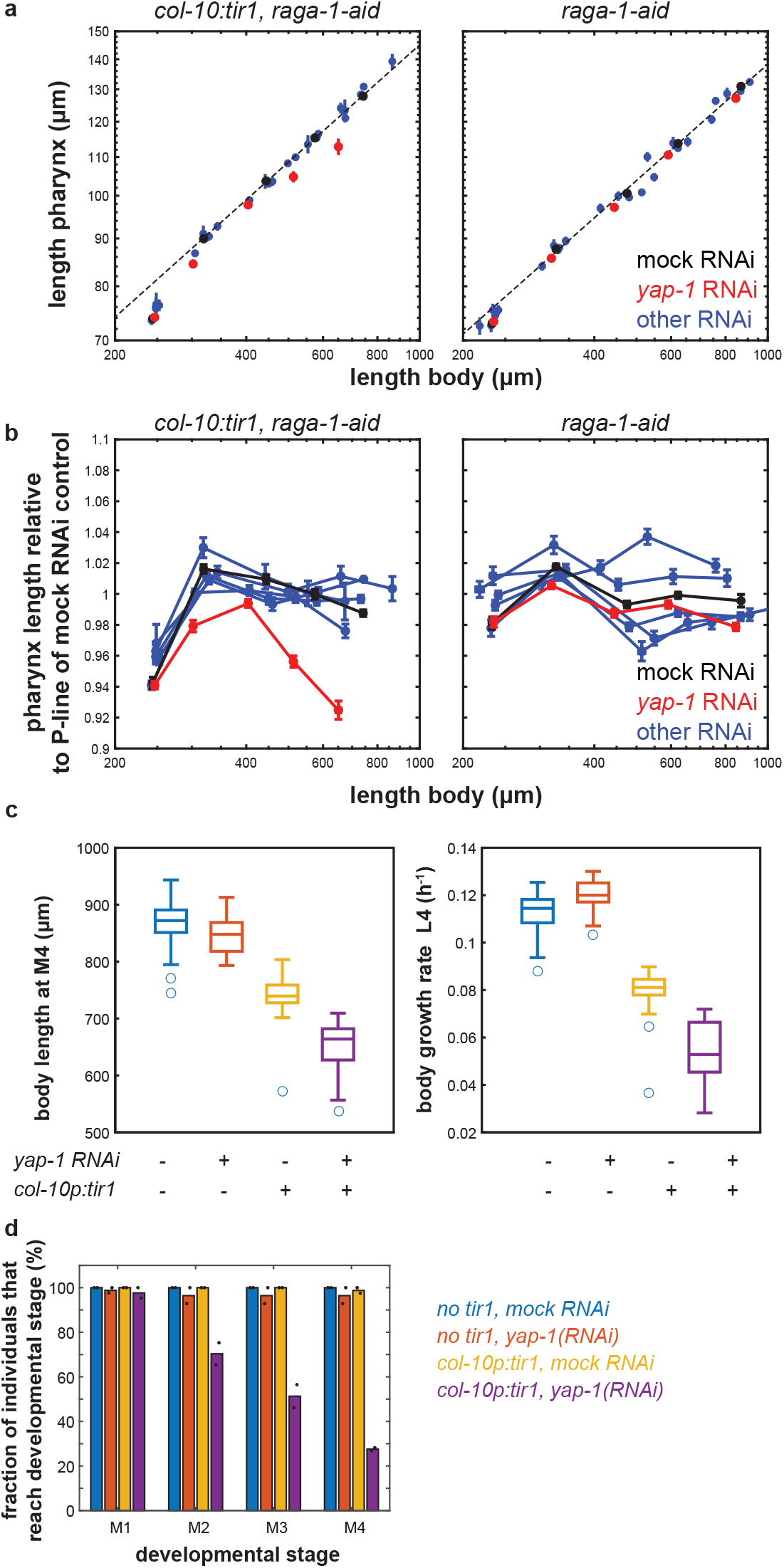
*yap-1* is required for robustness of pharynx-to-body length proportions towards epidermal growth inhibition. a. Pharynx vs. body length at 30% of L1 stage and at moults M1 to M4 under RAGA-1 AID in the epidermis (left) and without Tir1 expression (right) in combination with the indicated RNAi. Black: mock control, red: *yap-1*, blue: *dbl-1, lon-1, wts-1, ftt-2*. Black line: linear regression to M1 to M4 of mock RNAi. Error bars indicate standard error of the mean among individuals in x and y direction. If no error bars are visible, they are smaller than the marker size. See Supplemental Table 2 for details on number of individuals. b. as a., but for deviation of pharynx length from unperturbed conditions (deviation from P-line fitted to the respective mock RNAi control). p<10^−10^ (ranksum test) for all moults for comparison *yap-1(RNAi)* vs. mock RNAi for *col-10p:tir1* strain at M1 to M4. c. Box plot of growth rate and length at the 4th larval stage at indicated conditions measured in micro chambers. Scatter indicates individual animals. central line: median, box: interquartile ranges (IQR), whisker: ranges except extreme outliers (>1.5*IQR), circles: extreme outliers. p<10^−9^ (two-sided ttest) for all comparisons except unperturbed vs. *yap-1(RNAi)*. d. Fraction of individuals reaching the indicated developmental stage within the duration of the experiment. Black circles indicate values from 2 independent day-to-day repeats.

### Depletion of *yap-1* impairs larval development upon growth inhibition of the epidermis

To ask if the *yap-1* dependent response to tissue-selective depletion of RAGA-1 was physiologically important, we quantified the growth and development of *yap-1(RNAi)* animals in micro chambers. Consistent with previous reports ^34^, knock-down of *yap-1* alone did not impair development and fertility, and had only a minor impact on the overall growth rate (Fig. 5c). However, in combination with epidermal AID of RAGA-1, *yap-1(RNAi)* caused high penetrance larval arrest (Fig. 5d). This synthetic interaction between *yap-1* and epidermal RAGA-1 depletion also occurred on agar plates (Supplemental Fig. 6c). Larval arrest is therefore not caused by physical constriction of growth by the micro chambers. In conclusion, while *yap-1* is dispensable for development of *C. elegans* under standard laboratory conditions, *yap-1* is crucial under conditions of imbalanced tissue growth and for robustness of pharynx-to-body size proportions to tissue-specific perturbation of growth signalling.

## Discussion

Using time-lapse microscopy, we have observed near perfect size uniformity of the *C. elegans* pharynx with a coefficient of variation among individuals of less than 5%. The size heterogeneity of the *C. elegans* pharynx is thus substantially smaller than the heterogeneity in total body volume ^12^ (Fig. 1). Indeed, unlike total body volume size deviations, pharyngeal size deviations are corrected by sub-exponential growth such that large pharynxes undergo a smaller volume fold change than small pharynxes. This relation closely mimics a so-called adder mechanism, where the added volume per larval stage is independent of the volume at the beginning of the larval stage (Fig. 2). Adders were previously observed in bacteria, yeasts, and mammalian cells^7,9,36^. However, despite these phenomenological parallels, uni-cellular and multi-cellular adders are likely mechanistically distinct. Unlike cellular adders, which function cell autonomously, the systemic effects of tissue-specific growth inhibition suggest that pharyngeal size homeostasis involves communication among different tissues with each other (Fig. 3).

Since the pharynx is crucially involved in food uptake, deviations in its size may passively invoke a systemic effect on body growth. Food uptake is indeed likely to contribute to the robustness of pharyngeal size proportions, but multiple lines of evidence show that the robustness we observed involves additional mechanisms. First, endogenous fluctuations in the pharynx size were uncorrelated to overall body growth, suggesting that the pharynx size is not limiting for body growth within the endogenously observed range (Supplemental Fig. 2e). Second, we observe an ultra-sensitive response of body growth to pharynx size, where reducing pharynx length by only 5% caused a reduction of total body growth by more than 50%. This finding is difficult to reconcile with a mechanism based on food limitation alone. Indeed, simulations show that this ultra-sensitive response is required to produce the observed robustness of pharynx-to-body proportions to growth inhibition (Fig. 4). Third, we show that depletion of the gene *yap-1* impairs the robustness of pharynx-to-body proportions to tissue-specific growth inhibition. All these findings argue against a purely passive mechanism through food uptake and suggest a mechanism involving molecular regulation (Fig. 5).

In *C. elegans, yap-1* has previously been implicated in cell polarity^37,38^ and in the regulation of aging^34,39^. However, a role for *yap-1* in organ growth and size control of *C. elegans*, had not been characterized so far. We show a function of YAP-1 in organ growth coordination that becomes apparent under spatial imbalance of mTORC1 signalling. RNAi of other genes of the Hippo pathway did not impair pharynx-to-body size proportions (*ftt-2*/14-3-3 and *wts-1/H*ippo kinase). However, based on the function of their human orthologs, these genes are likely negative regulators of YAP-1, such that their genetic depletion would hyperactivate *yap-1* ^34^ rather than phenocopy its depletion. Future experiments will differentiate Hippo kinase dependent and independent regulation of YAP-1 in *C. elegans*.

Consistent with our results, the *Drosophila* homolog of YAP-1 (called Yorkie) transcriptionally controls the relaxin-like hormone *dilp8*, a systemic growth regulator of *Drosophila* larvae ^40^. *C. elegans* YAP-1 may similarly act upstream of hormonal control, or directly control the transcription of growth-related genes. Fly and mammalian YAP-1 orthologs respond to a variety of mechanical stimuli. Since the gastro-intestinal tract of *C. elegans* is attached to the exoskeletal cuticle, changes in pharyngeal length are likely to create mechanical forces between cells and tissues, or between cells and the cuticle. Thus, an attractive model for the ultra-sensitive coupling between pharynx and body growth is that marginal disproportions between tissues create pulling forces between them, leading to YAP-1 regulation. An important open question is how cells quantitatively convert such forces into an appropriate response in gene expression and growth, which can now be addressed in the context of an entire living animal.

## Materials and Methods

### List of strains used in this study

All strains used in this study were created by genetic crosses from the following previously published alleles: *bqSi577* ^41^, *wbmIs88* ^42^, *ieSi60* ^23^, *daf-2(bch40)* ^43^, *raga-1(wbm40)* ^26^, *reSi1* ^27^, *xeSi376* ^23^, *daf-16(mu86)* ^44^.

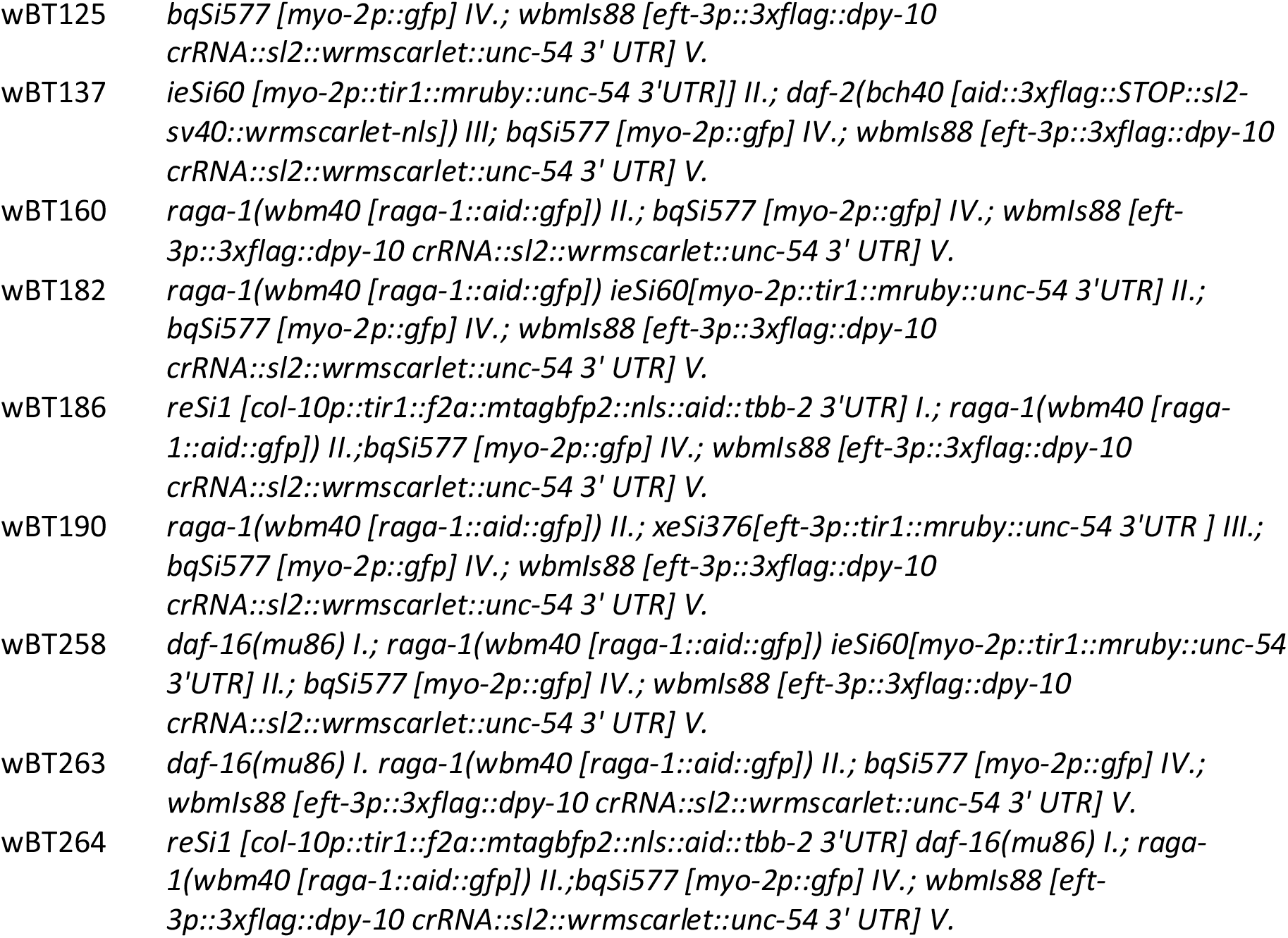

### Micro chamber preparation

Micro chambers were manufactured as described in ^12^ using a master PDMS mould as a micro comb to produce wells in a 4.5% Agarose gel in S-basal. For all experiments, chamber dimensions were 600×600×20μm. As a food source, the bacterial strain OP50-1 was grown on NGM plates by standard methods, scraped off using a piece of agar and then filled into the wells of the agarose gel. Wells were filled with eggs at 2-fold stage and subsequently inverted onto a dish of 3.5cm diameter with high optical quality gas-permeable polymer bottom (ibidi). The remaining surface of the dish was covered with 3% low melting temperature Agarose dissolved in S-basal (cooled down to below 42 °C prior to application). The agarose was overlayed with ^~^0.5ml polydimethylsiloxane (PDMS) and the dish was sealed with parafilm to prevent excessive evaporation. PDMS was allowed to cure at room temperature on the microscope during the acquisition. Using a custom-made plate holder, six dishes could be imaged simultaneously on one microscope. Auxin (IAA, Sigma) solutions were freshly prepared on the day of the experiment as a 400x stock in EtOH and subsequently diluted to the indicated concentration in agarose to a final EtOH concentration of 0.25% immediately prior to use for micro chamber assembly.

### Imaging

All experiments were performed on a Nikon Ti2 epifluorescence microscope using a 10x objective with NA=0.45 and a Hamamatsu ORCA Flash 4 sCMOS camera. Temperature was maintained at 25 +/-0.1 °C by incubator enclosing the entire microscope and a feedback controlled temperature regulator (ICE cube, life imaging services). Separately triggerable LEDs (SpectraX, lumencore) for 491nm (GFP) and for 575nm (mCherry/mScarlet) were used as an excitation light source that was TTL-triggered to ensure rapid switching between wavelengths and thus imaging of red and green fluorescence within 10ms. Acquisition times were kept below 10ms, such that two color imaging could be completed within 30ms. Rapid image registration was crucial to minimize a shift between fluorescent channels in the absence of physical or chemical immobilization of the animals. Software autofocus as provided by Nikon’s NIS software was used every 10 minutes using 575nm excitation at low intensity. We confirmed that these acquisition settings did not impair growth, development, and fertility of animals.

### Image analysis

A custom-made Matlab script was used to segment worms and pharynxes from raw fluorescent images. For treatment of the total body of worms, all procedures were identical to those previously described ^12^. For the pharynx, a pixel classifier was trained using Ilastik ^45^ on the green fluorescent channel. Segmentations of body and pharynx were subsequently straightened ^12^ using the body outline as a template. Pharyngeal and body volume and length were computed from these straightened images assuming rotational symmetry ^22^. A decision tree based classifier ^12^ was used to determine the time point of hatching, and to identify time points where straightening failed (e.g. due to self-touching animals). Detection of moults and computation of growth rates was conducted as previously described using a custom-made Matlab script ^12^. Trendlines in scatter plots were computed using robust linear regression using the robustfit() method of Matlab (v2021b) and default parameters. We thereby reduce sensitivity to outliers.

### RNAi

All RNAi experiments were conducted by feeding using HTT115 clones retrieved from available libraries ^46,47^ and validated by Sanger sequencing. (clone numbers: *dbl-1*: sjj_T25F10.2; *lon-1*: sjj_F48E8.1, *yap-1*: sjj_F13E6.4, *ftt-2*: sjj_F52D10.3, *wts-1*: sjj_T20F10.1, *rsks-1*: sjj2_Y47D3A.16). RNAi was initiated on plates one generation prior to loading in micro chambers. For RNAi plate preparation, bacterial clones were grown to saturation over night in LB with Ampicillin (100μg/ml) and dispensed on NGM plates NGM plates containing 1mM IPTG and 50μg/ml Carbenicillin. L4 stage animals grown on OP50-1 were transferred to RNAi plates and their embryonic progeny was transferred into micro chambers containing bacteria expressing the same RNAi clone using an eyelash glued to a pipette tip. For RNAi inside micro chambers, bacterial over night cultures were induced in liquid for 4h using 4mM IPTG, concentrated by centrifugation, and dispensed onto NGM plates containing 1mM IPTG and 50μg/ml Carbenicillin. After drying, bacteria were scraped off the plate and used as described above for OP50-1.

### Comparison of observed growth to randomized simulations

Simulations were performed as described ^12^ and started with the measured distribution of pharynx volume after hatching and each individual was assigned one of the following parameters randomly selected from the experimentally measured distributions, depending on the simulation: i) a pharyngeal growth rate and a larval stage duration (independently randomized), ii) the volume fold change, iii) the volume added within one larval stage. From these randomly selected parameters the pharyngeal volume of the next larval stage was computed and the process was iterated until the end of L4. Each randomization was repeated 1000 times for each day-to-day repeat and the pooled for visualization. To exclude effects of large heterogeneity in pharyngeal volume at hatch, randomizations were also started at the end of L1 instead of at hatching, which yielded equivalent conclusions.

### Averaging of growth rates and volumes after re-scaling

To average trajectories of growth rates, volume, and length without perturbing alignment of moults individual trajectories were re-scaled to the larval stage duration and averaged after interpolation at 100 points per larval stage. Data was then averaged over all individuals at each of the 100 points per larval stage, and the averaged data was rescaled to the average larval stage duration.

### Computation of width and growth rates

Unless indicated otherwise, average growth rates per larval stage were calculated as the difference between the natural logarithm of the volume at the beginning and the end of the larval stage divided by the larval stage duration (μ = Δlog(V)/Δt). Average width (*w*) of body and pharynx was calculated from volume *v* and length *l* as follows: 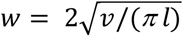.

## Author contributions

BDT and KS conceived the study, all experiments except those mentioned below were executed by KS. Experiments in Figure 1 were conducted by BDT, experiments in Figure 6c were conducted by IG. BDT and KS analysed the data. BDT designed and analysed the mathematical model. WBM and AL created and validated the *raga-1-aid* allele. KS and IG created the strains used for the study. BDT wrote the manuscript. KS and WMB edited the manuscript.

## Acknowledgements

We are thankful to Cihan Elci for technical assistance and to Rutger Hermsen and Helge Grosshans for useful discussions and helpful feedback on the manuscript. We acknowledge support by the Microscopy Imaging Center at the University of Bern. This work received funding from the Swiss National Science Foundation (SNSF) in the form of an Eccellenza Professorial Fellowship (PCEFP3_181204) to B.D.T. and from the Novartis Foundation for Medical-Biological Research (Grant #20A011) Some strains were provided by the CGC, which is funded by NIH Office of Research Infrastructure Programs (P40 OD010440).

