## Supplemental Figures and Tables for "Ultra-sensitive coupling between organ growth and size by YAP-1 ensures uniform body plan proportions in *C. elegans*"

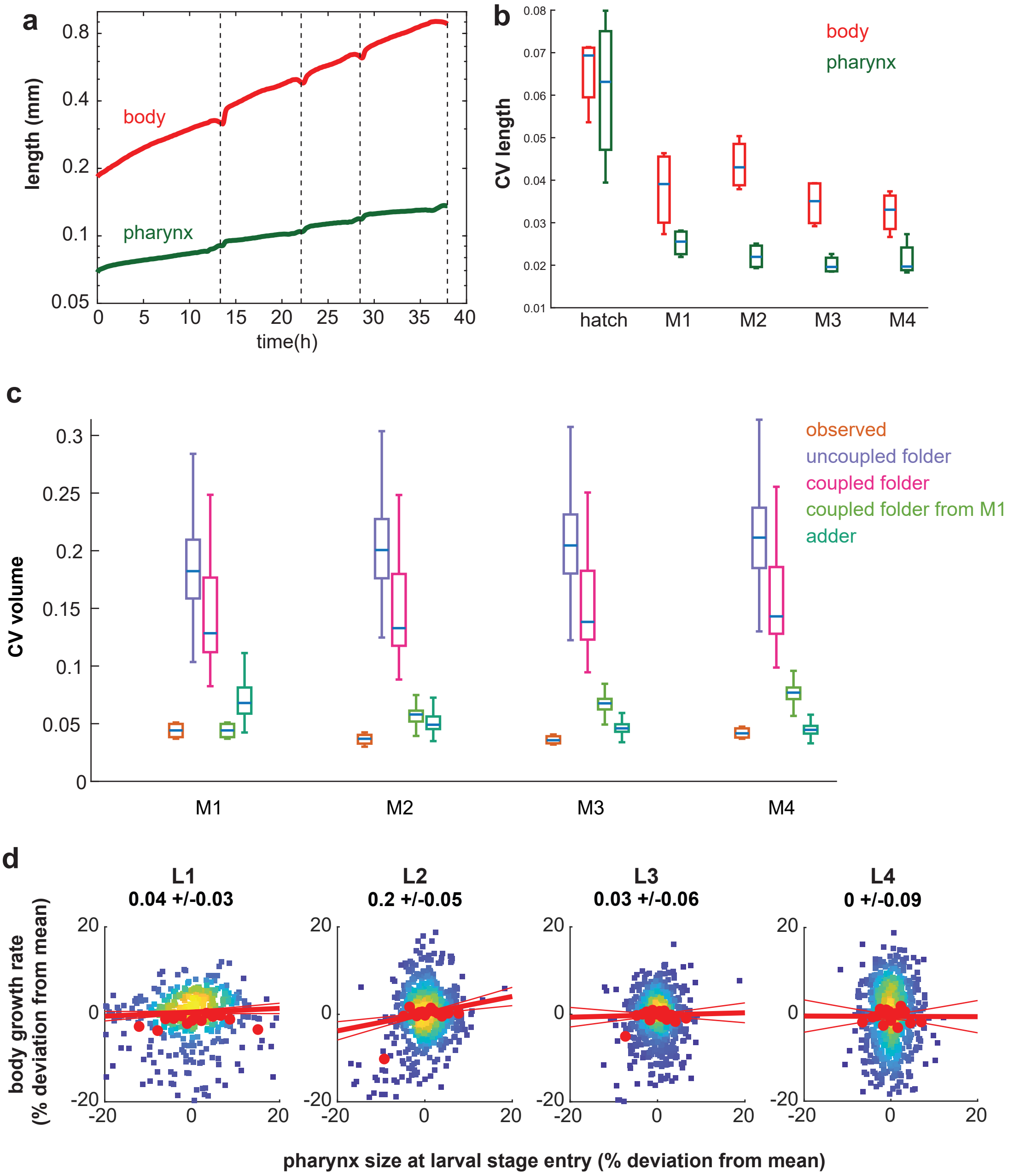


Supplemental Figure 1

**Supplemental Figure 1. Pharyngeal length is less heterogeneous among individuals than total body length**

a. Body (red) and pharynx (green) length as a function of time averaged from n=475 individuals. For averaging, individuals of each larval stage were re-scaled to have matching larval stage entry and exit points and the growth curve was scaled back to the mean larval stage duration.
b. Coefficient of variation (CV) of body (red) and pharynx (green) length at hatch and larval moults. Box plots represent CVs of 4 independent day-to-day repeats with $105 < n < 181$ individuals per repeat.
c. Comparison of observed CV of pharyngeal volume to randomized simulations at indicated larval moults.
d. Scatter plot of pharynx volume at larval stage entry vs. body growth rate for individual animals at indicated larval stages. Red circles: binned average along x-Axis. red line: robust linear regression (thick) with 95% confidence intervals (thin). Data is shown as relative deviation to the batch mean. Rare outliers beyond the +/- 20% range are omitted for clarity. Value above chart indicates slope of regression line +/- 95% CI

Boxplots in b,c: central line: median, box: interquartile ranges (IQR), whisker: ranges except extreme outliers (>1.5*IQR)

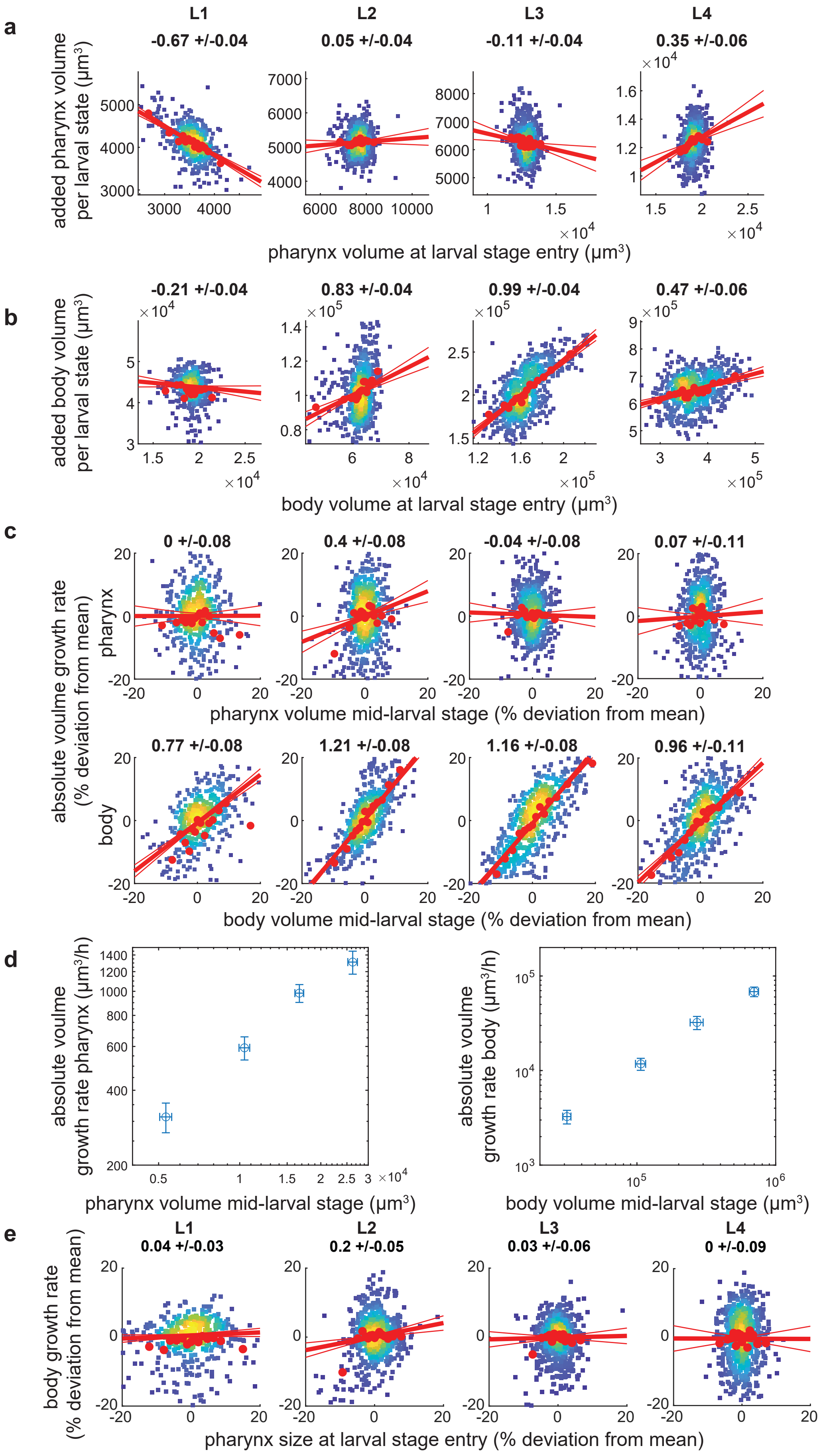


Supplemental Figure 2

**Supplemental Figure 2. Linear pharyngeal volume growth within larval stages produces and adder-like behaviour**

a. Scatter plot of pharynx volume at larval stage entry vs. added pharynx volume per larval stage for each individual. Numbers above chart indicate slope of regression line +/- confidence interval. Red circles: binned average along x-Axis. red line: robust linear regression (thick) with confidence intervals (thin). n = 475,641, 641, 638 individuals for L1 to L4 from 4 day-to-day repeats.
b. As a., but for body volume
c. As a., but for pharyngeal and body volume at 40% of the larval stage vs. absolute rate of pharynx and body volume increase. Data was normalized to batch mean. Outliers outside of the +/- 20% range are omitted for better visualization.
d. Mean pharyngeal and body volume at 40% of the larval stage (L1 to L4 from left to right) vs. absolute rate of volume increase. Error bars: standard deviation. Near linear scaling of growth and size in a log-log plot is consistent with near exponential growth across larval stages. Number of individuals n as in a-c.
e. As a., but for pharynx volume at larval stage entry vs. body volume growth rate ($\Delta$log(V)/$\Delta$t) in the same larval stage.

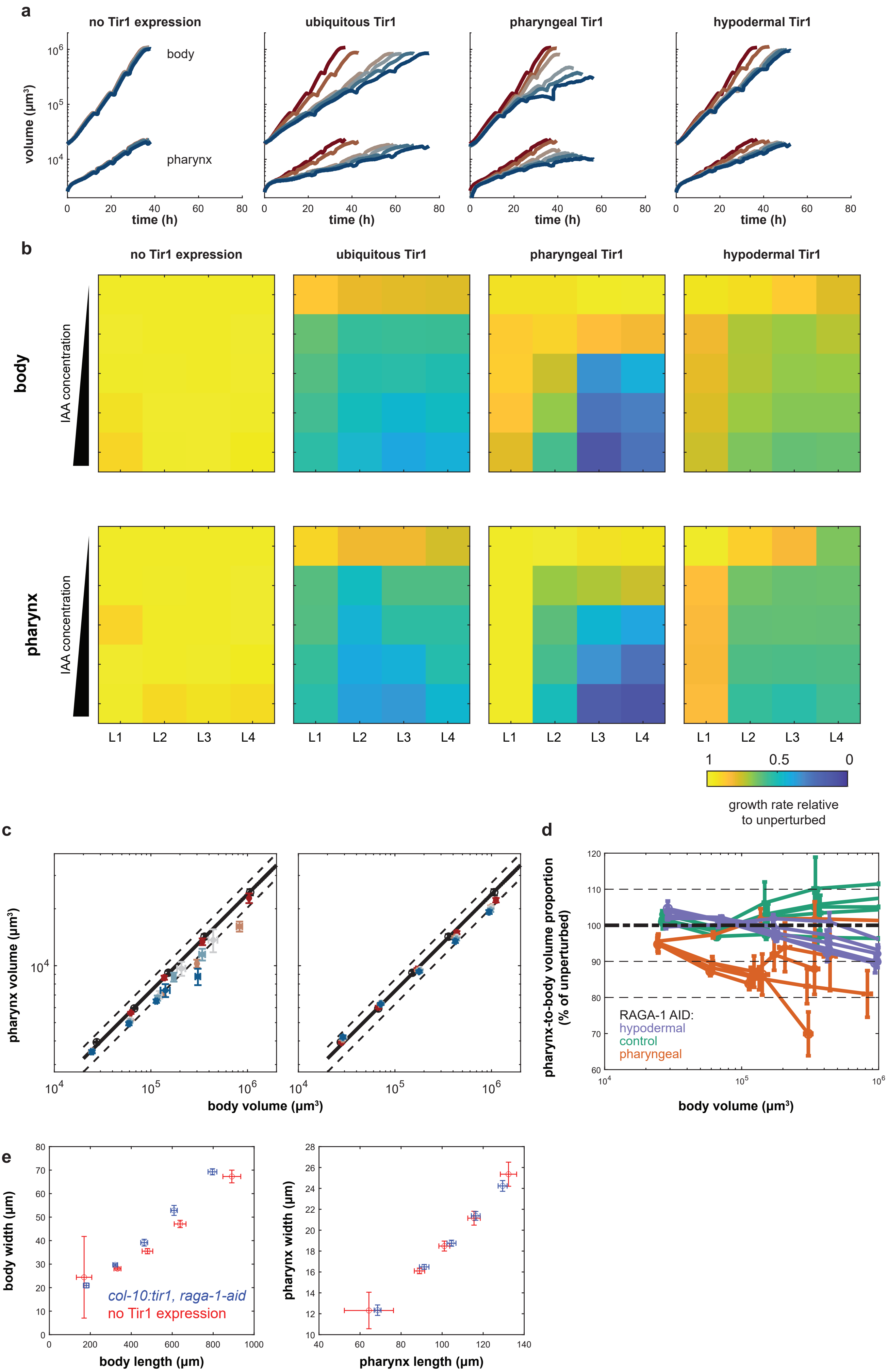


Supplemental Figure 3

**Supplemental Figure 3. Pharynx-to-body volume proportions are robust to tissue-specific depletion of RAGA-1**

a. Body (top) and pharynx (bottom) volume as a function of time with depletion of RAGA-1 in indicated tissues. Color indicates IAA concentration from red to blue: no Tir1 expression 0mM IAA, Tir1 expression + 0mM, 0.1mM, 0.25mM, 0.5mM, 1mM IAA. Number of individuals n, 26 < n < 259 from number of day-to-day repeats m, 3 < m < 11. See Supplemental Tables 1 and 2 for n and m of each condition.
b. Heatmap showing volume growth rate of body and pharynx normalized to unperturbed growth (no Tir expression and no IAA) after RAGA-1 depletion by AID in indicated tissues, larval stages, and IAA concentrations. IAA concentrations increase from top to bottom as follows: 0mM, 0.1mM, 0.25mM, 0.5mM, 1mM
c. Scatter plot showing body vs. pharynx volume at the beginning of L1 and at all larval moults M1 to M4 (circles in order from left to right) under pharyngeal (left) or epidermal (right) AID of RAGA-1. black circles: relation between pharynx and body length when unperturbed (no Tir expression and no IAA). Solid black line: linear regression to unperturbed body-to-pharynx length (P-line). Dashed black line: 15% deviation from P-line. Coloured circles: IAA concentrations from red to blue: 0, 100, 250, 500, 1000µM. Error bars are standard error of the mean among day-to-day repeats.
d. Deviation of pharynx-to-body volume ratio from the unperturbed P-line vs. body volume at early L1 and larval moults. Colours indicate different Tir1 expression; orange: pharynx, purple: epidermis, green: no Tir1 expression. Circle size indicates IAA concentration from 0mM (smallest circle) to 1mM (largest circle) as indicated in c. Error bars are standard errors among day-to-day repeats.
e. Body and pharynx length vs. width for *col10p:tir1; raga-1-aid* (blue) and for *raga-1-aid* without Tir1 expression (red) at 1mM IAA at hatch and all four moults from left to right. Error bars are the standard deviation among individual animals.

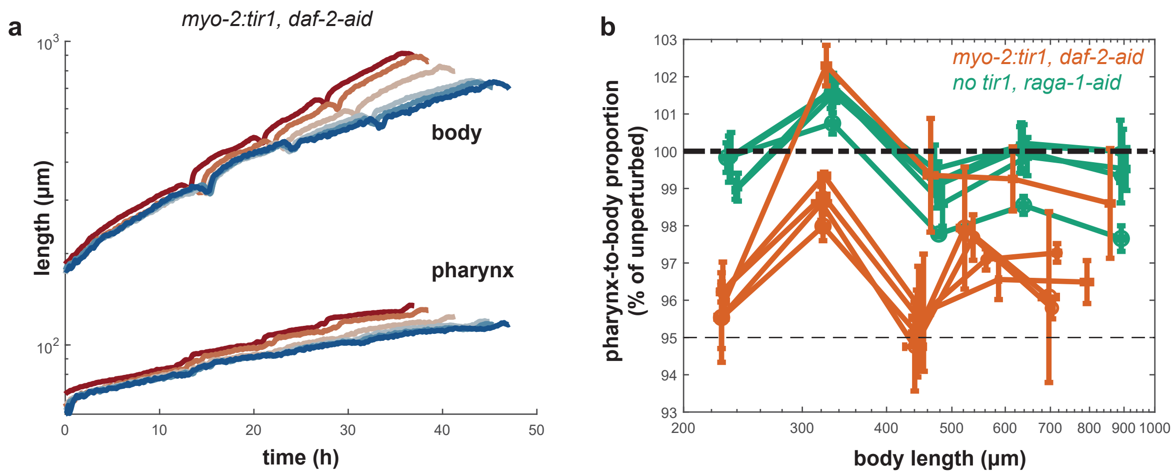


Supplemental Figure 4

**Supplemental Figure 4. Pharynx-to-body volume proportions are robust to tissue-specific depletion of DAF-2**

a. Body (top) and pharynx (bottom) length as a function of time with depletion of DAF-2 in the pharynx. Color indicates IAA concentration from red to blue as follows: no Tir1 expression 0mM IAA, Tir1 expression + 0mM, 0.1mM, 0.25mM, 0.5mM, 1mM IAA.

b. Deviation from unperturbed pharynx-to-body length ratio (deviation from P-line) vs. body length in early L1 and at larval moults upon DAF-2 AID in the pharynx (orange) and without DAF-2 AID (green). Circle size indicates IAA concentration from 0mM (smallest circle) to 1mM (largest circle) as indicated in c. Error bars are standard errors among day-to-day repeats. Thick dashed lines indicate unperturbed state (P-line). Thin dashed line indicates a 5% deviation from te P-line.

a,b. 17 < n <58 individuals from 2 day-to-day repeats. See Supplemental Tables 1 and 2 for exact number of individuals for each condition

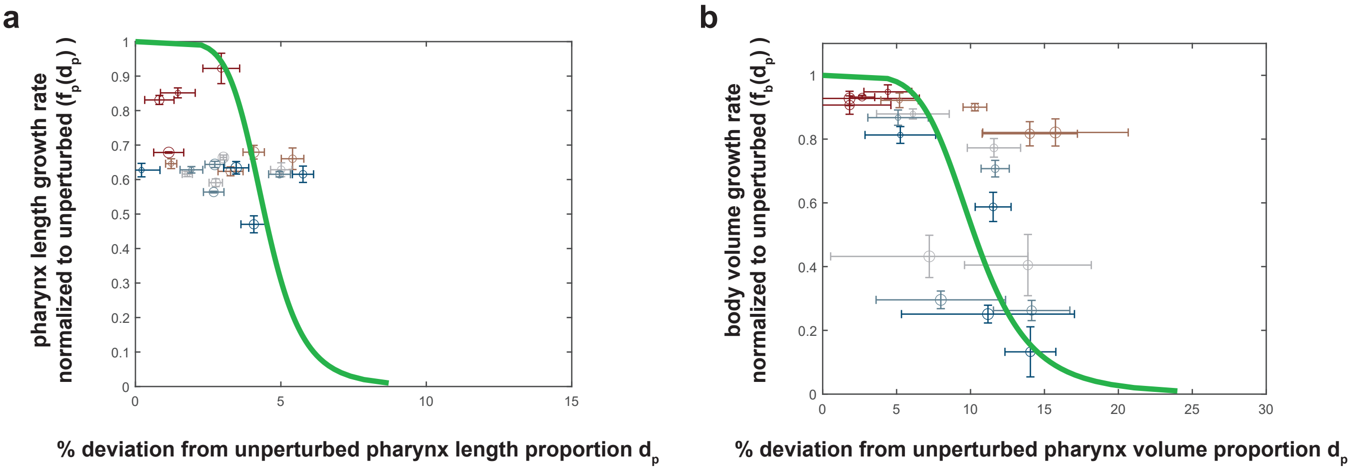


Supplemental Figure 5

**Supplemental Figure 5. An ultra-sensitive relation between pharynx size and body growth is required for robustness of pharynx-to-body length proportions to tissue-specific growth inhibition**

a. Pharynx length growth rate as a function of deviation of pharynx length from the P-line under epidermal growth inhibition normalized to unperturbed growth. Colour indicates IAA concentration increasing from red (0 mM) to blue (1 mM). Circle size indicates the larval stage. Green line is the Hill function obtained from fit to the pharyngeal RAGA-1 AID described in the main text and shown in Figure 4a.
b. Body volume growth rate as a function of deviation of pharynx volume from the P-line under pharyngeal growth inhibition normalized to unperturbed growth. Colour indicates IAA concentration increasing from red (0 mM) to blue (1 mM). Circle size indicates the larval stage. Green line is a fitted Hill function as described in the main text.

a,b. error bars indicate standard error of the mean between day-to-day repeats.

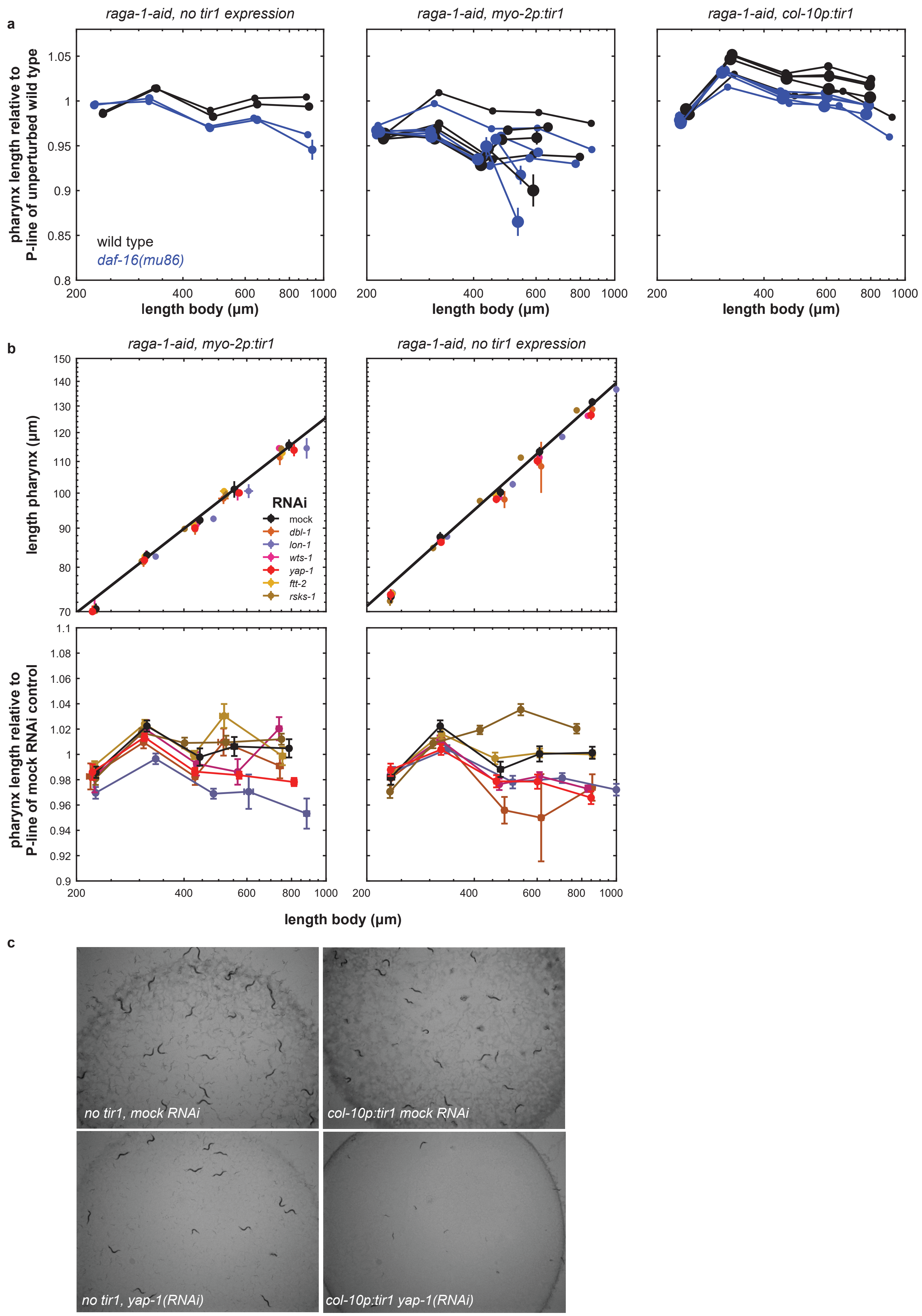


Supplemental Figure 6

**Supplemental Figure 6. *yap-1* is required for robustness of pharynx-to-body volume proportions to epidermal growth inhibition**

a. Deviation of pharyngeal length of *daf-16(mu86)* mutants from P-line fitted to the unperturbed wild type. Connected circles indicate 30% L1, and M1 to M4 moults (left to right) of indicated genotypes and RAGA-1 AID treatments. IAA concentrations: without Tir1 expression (left): 0mM, 0.1mM. For pharyngeal (middle) and epidermal (right) Tir1 expression: 0mM, 0.1mM, 0.25mM, 0.5mM, 1mM. Marker size indicates IAA concentration (smallest marker for lowest concentration). Error bars are standard error of the mean among individuals. See Supplemental Tables 1 and 2 for number of individuals and day-to-day repeats of each condition.
b. Top: body length vs. pharynx length for *raga-1-aid* strain with Tir1 expression in the pharynx (left) or without Tir1 expression (right) for indicated RNAi. Bottom: body length vs. deviation of pharyngeal length from P-line fitted to the mock RNAi control for indicated RNAi. IAA concentration: 0.1mM. error bars are standard error of the mean among individuals.
c. Images of animals grown on RNAi plates under indicated conditions. RNAi was initiated from the L4 stage of the parental generation. Animals were synchronized at L1 stage by bleaching and grown on RNAi plates containing 0.5mM IAA for 72 hours.

**Supplemental Table 1. number of day-to-day repeats per condition**

| IAA | no Tir1 expression, *raga-1-aid* | *eft-3p:tir1, raga-1-aid* |
|---|---|---|
| 0mM | 10 | 4 |
| 0.1mM | 10 | 4 |
| 0.25mM | 7 | 4 |
| 0.5mM | 7 | 4 |
| 1mM | 7 | 4 |

| IAA | *myo-2p:tir1, raga-1-aid* | *col-10p:tir1, raga-1-aid* |
|---|---|---|
| 0mM | 8 | 4 |
| 0.1mM | 8 | 4 |
| 0.25mM | 5 | 4 |
| 0.5mM | 5 | 4 |
| 1mM | 5 | 4 |

| IAA | *myo-2p:tir-1, daf-2-aid* | *myo-2p:tir1, raga-1-aid, daf-16(mu86)* |
|---|---|---|
| 0mM | 2 | 5 |
| 0.1mM | 2 | 5 |
| 0.25mM | 2 | 2 |
| 0.5mM | 2 | 2 |
| 1mM | 2 | 2 |

| IAA | *col-10p:tir1, raga-1-aid, daf-16(mu86)* | no Tir1 expression, *raga-1-aid*, *daf-16(mu86)* |
|---|---|---|
| 0mM | 2 | 2 |
| 0.1mM | 2 | 2 |
| 0.25mM | 2 | na |
| 0.5mM | 2 | na |
| 1mM | 2 | na |

**Supplemental Table 2. total number of individuals tested**

no Tir1 expression, *raga-1-aid*

| IAA | L1 | L2 | L3 | L4 |
|---|---|---|---|---|
| 0mM | 142 | 244 | 242 | 220 |
| 0.1mM | 104 | 171 | 171 | 153 |
| 0.25mM | 53 | 107 | 106 | 98 |
| 0.5mM | 61 | 98 | 98 | 89 |
| 1mM | 83 | 123 | 120 | 99 |

*eft-3p:tir1, raga-1-aid*

| IAA | L1 | L2 | L3 | L4 |
|---|---|---|---|---|
| 0mM | 52 | 68 | 68 | 66 |
| 0.1mM | 53 | 86 | 86 | 84 |
| 0.25mM | 52 | 71 | 69 | 61 |
| 0.5mM | 50 | 75 | 72 | 58 |
| 1mM | 55 | 80 | 71 | 57 |

*myo-2p:tir1, raga-1-aid*

| IAA | L1 | L2 | L3 | L4 |
|---|---|---|---|---|
| 0mM | 159 | 258 | 258 | 236 |
| 0.1mM | 135 | 191 | 189 | 142 |
| 0.25mM | 81 | 114 | 114 | 88 |
| 0.5mM | 46 | 90 | 86 | 65 |
| 1mM | 56 | 95 | 90 | 33 |

*col-10p:tir1, raga-1-aid*

| IAA | L1 | L2 | L3 | L4 |
|---|---|---|---|---|
| 0mM | 77 | 106 | 105 | 105 |
| 0.1mM | 41 | 70 | 70 | 69 |
| 0.25mM | 70 | 103 | 103 | 101 |
| 0.5mM | 58 | 93 | 92 | 91 |
| 1mM | 27 | 42 | 42 | 39 |

*myo-2p:tir-1, daf-2-aid*

| IAA | L1 | L2 | L3 | L4 |
|---|---|---|---|---|
| 0mM | 27 | 47 | 47 | 47 |
| 0.1mM | 34 | 49 | 49 | 49 |
| 0.25mM | 32 | 57 | 57 | 57 |
| 0.5mM | 24 | 41 | 41 | 39 |
| 1mM | 18 | 31 | 31 | 29 |

*myo-2p:tir1, raga-1-aid, daf-16(mu86)*

| IAA | L1 | L2 | L3 | L4 |
|---|---|---|---|---|
| 0mM | 105 | 152 | 151 | 134 |
| 0.1mM | 154 | 198 | 195 | 140 |
| 0.25mM | 64 | 89 | 86 | 71 |
| 0.5mM | 32 | 51 | 51 | 42 |
| 1mM | 41 | 71 | 71 | 38 |

*col-10p:tir1, raga-1-aid, daf-16(mu86)*

| IAA | L1 | L2 | L3 | L4 |
|---|---|---|---|---|
| 0mM | 34 | 61 | 61 | 59 |
| 0.1mM | 23 | 40 | 40 | 39 |
| 0.25mM | 39 | 57 | 57 | 57 |
| 0.5mM | 25 | 41 | 41 | 40 |
| 1mM | 36 | 45 | 45 | 44 |

no Tir1 expression, *raga-1-aid*, *daf-16(mu86)*

| IAA | L1 | L2 | L3 | L4 |
|---|---|---|---|---|
| 0mM | 50 | 102 | 101 | 84 |
| 0.1mM | 48 | 77 | 77 | 57 |

no Tir expression, raga-1-aid, 0.5 mM IAA, mock RNAi

| hatch | M1 | M2 | M3 | M4 |
|---|---|---|---|---|
| 39 | 72 | 71 | 71 | 66 |

no Tir expression, *raga-1-aid*, 0.5 mM IAA, *yap-1(RNAi)*

| hatch | M1 | M2 | M3 | M4 |
|---|---|---|---|---|
| 62 | 71 | 70 | 70 | 70 |

*col-10p:tir1*, *raga-1-aid*, 0.5 mM IAA, mock RNAi

| hatch | M1 | M2 | M3 | M4 |
|---|---|---|---|---|
| 36 | 63 | 63 | 63 | 62 |

*col-10p:tir1*, *raga-1-aid*, 0.5 mM IAA, *yap-1(RNAi)*

| hatch | M1 | M2 | M3 | M4 |
|---|---|---|---|---|
| 65 | 98 | 80 | 59 | 31 |
